# Glucocorticoid and Mineralocorticoid Receptors Jointly Promote Vascular Development in Kidney Organoids

**DOI:** 10.1101/2025.09.03.673963

**Authors:** Cory P. Johnson, Hannah M. Somers, Sophie E. Craig, Heath Fuqua, Lynne Beverly-Staggs, Kailee E. Tanaka, Sydney M. Brown, Charles H. Toulmin, Matthew D. Cox, Joel H. Graber, Melissa S. Maginnis, Hermann Haller

**Affiliations:** Kathryn W. Davis Center for Regenerative Biology and Aging, MDI Biological Laboratory, Bar Harbor, ME, USA; Department of Nephrology, Hannover Medical School, Hannover, Germany; Graduate School of Biomedical Science and Engineering, University of Maine, Orono, Maine, USA; Department of Molecular and Biomedical Sciences, University of Maine, Orono, Maine, USA

## Abstract

To examine the co-development of vasculature and renal epithelial tissue, we employed a human pluripotent stem cell-derived kidney organoid system. We found that cooperative signaling through the glucocorticoid receptor and mineralocorticoid receptor via hydrocortisone enabled rich endothelial cell differentiation and vessel formation. Bulk RNA sequencing analysis revealed that hydrocortisone perturbs an angiogenic transcriptional program early in development and promotes instead a pro-endothelial survival transcriptional program, with upregulation of angiopoietin 1 at both the mRNA and protein level. Additionally, we saw that hydrocortisone does not seem to significantly affect gene expression of canonical nephrogenic genes compared to our controls, suggesting its effect is largely restricted to endothelial cell differentiation. Our results show that kidney organoids offer a unique platform to study developmental signals that drive endothelial cell differentiation and vessel formation.

## Introduction

Organogenesis requires precise signal coordination and cooperation among: a) stromal cells, i.e., connective-tissue cells present in any organ, such as fibroblasts and endothelial cells (ECs) (Roberts et al., 2016); and b) parenchymal cells, i.e., organ-specific cells such as epithelial cells. During organogenesis, ECs directly influence the growth and differentiation of the parenchyma, and, in turn, parenchymal cells directly influence the development and maintenance of endothelial networks, with strict spatial and temporal signaling required in each direction (Ribatti et al., 2023). To investigate this complex coordination, we must develop tools that accurately model the development of both the target organ and the vasculature simultaneously.

Kidney organoids have revolutionized the study of human kidney development. Recent advances in protocols for kidney organoid differentiation have significantly improved nephron maturation and function (Abdollahzadeh et al., 2024; Aceves et al., 2022; Clerkin et al., 2025; Huang et al., 2024; Kearney et al., 2025; Kim et al., 2022; Krupa et al., 2024; Kuang et al., 2025; Nerger et al., 2024; Ruiter et al., 2022; Sarami et al., 2025; Vanslambrouck et al., 2023; Wilson et al., 2025). And while other kidney organoid protocols have incorporated the addition of ECs and have even perfused the organoids with microfluidic vasculature (Garreta et al., 2024; Kroll et al., 2024; Maggiore et al., 2024), the understanding of the signals necessary for coordinating endothelial and nephrogenic differentiation in kidney organoids remains incomplete.

Hydrocortisone (HC), a glucocorticoid, has several significant roles in embryonic development including cell differentiation (Bridges et al., 2020; Nesan et al., 2012), proliferation (Crossin et al., 1997; Durant et al., 1986; Grisé et al., 2021), and metabolism (Vegiopoulos and Herzig, 2007). HC influences these biological processes by acting through the glucocorticoid receptor (GR) and the mineralocorticoid receptor (MR), which are known regulators of many developmental programs (Bridges *et al*., 2020; Martinerie et al., 2013; Nesan *et al*., 2012), including EC factors (Srivastava et al., 2019). We hypothesized that strategic timing of HC treatment during kidney organoid development would enhance EC differentiation and vessel formation. To test this, we aimed to first optimize HC treatment timing, then identify GR- vs. MR-specific contributions mediating HC-induced outcomes, and, finally, characterize the transcriptional responses guiding these processes. We show that HC indeed enhanced endothelial differentiation and vascular complexity; both GR and MR activation are required. We found that HC suppresses transcriptional expression of angiogenic genes during treatment, while enhancing the expression of the EC-survival factor, angiopoietin 1 (ANG1). Finally, HC preserved nephrogenic identity in both early and later developmental stages. This observation not only highlights a pivotal role for glucocorticoid signaling in vascular development but also provides a platform to study the impacts of other cell fate regulators in vascular and nephron development.

## Results

### Hydrocortisone Enhances Endothelial Differentiation and Vascular Formation

To evaluate the impact of HC on EC differentiation in kidney organoids, we first tested three HC treatment schemes (HC-A, HC-B, and HC-C; Fig. 1A), using a protocol adapted from the Bonventre lab (Morizane et al., 2015) and the Little lab (Kumar et al., 2019). Western blot analysis of day 28 organoids revealed robust upregulation of the EC markers CD31 and VE-cadherin (VECAD) in HC-A-treated groups compared to DMSO controls (CON), but not in HC-B- or HC-C-treated groups (Fig. 1B). Densitometry quantification confirmed increased CD31 and VECAD protein levels under HC-A treatment (Fig. 1C). Importantly, epithelial cell identity was preserved following HC-A treatment, as qPCR analysis at day 24 showed no significant downregulation of nephron segment markers cadherin 1 (ECAD; tubule) and nephrin (NPHS1; podocytes) (Fig. 1D and Supplemental Fig. 1).

**Figure 1:**
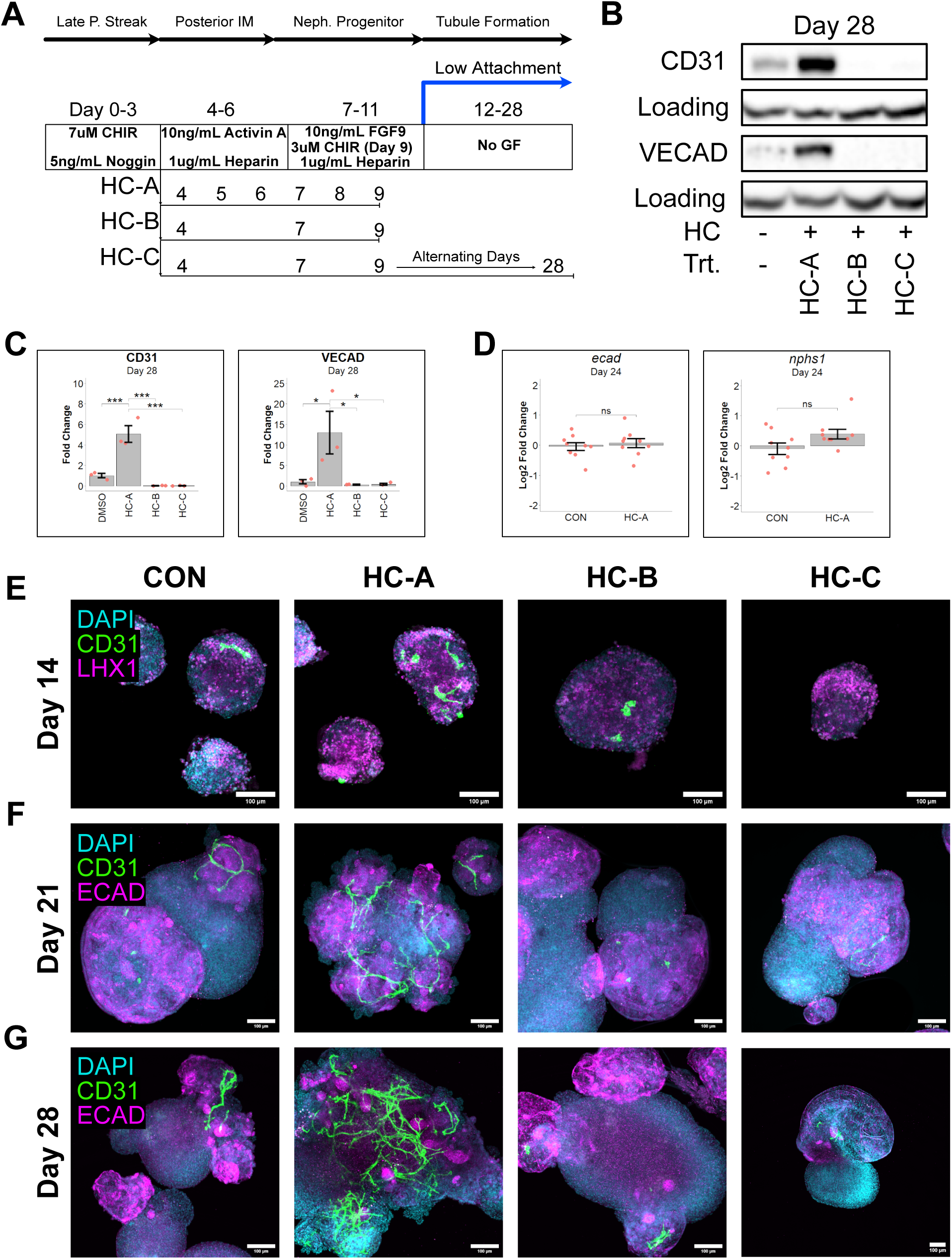
Optimization of HC-induced EC differentiation and vessel formation in kidney organoids. (A) Schematic of differentiation and HC treatment schemes tested. (B) Representative western blot for EC protein expression (CD31 and VECAD) in day 28 renal organoid cultures. (C) Densitometry quantification of CD31 (left panel) and VECAD (right panel) western blotting from panel B. (D) Log2 fold change of mRNA expression (HC-A-treated vs DMSO) of marker genes of distal tubule (*ecad*; left panel) and podocytes (*nphs1*; right panel) in 24-day-old kidney organoids. (E) Representative images on day 14 of DAPI (cyan), CD31 (green), and LHX1 (magenta) in DMSO-(CON; left panel), HC-A-(left-middle panel), HC-B-(right-middle panel) and HC-C-(right panel) treated organoids. (F) Representative images on day 21 of DAPI (cyan), CD31 (green), and ECAD (magenta) in DMSO-(CON; left panel), HC-A-(left-middle panel), HC-B (right-middle panel), and HC-C-(right panel) treated organoids. (G) Representative images on day 28 of DAPI (cyan), CD31 (green), and ECAD (magenta) in DMSO-(CON; left panel), HC-A-(left-middle panel), HC-B-(right-middle panel), and HC-C-(right panel) treated organoids. *n=3* biological replicates per treatment group (C only). *n=9* biological replicates combined across 3 experiments per treatment group (D only). Data are represented as mean ± SEM. * p< 0.05, ** p < 0.01, *** p < 0.001, and **** p < 0.0001.

Immunofluorescence analysis across three timepoints revealed progressive expansion of CD31+ vascular structures in HC-A treated organoids. We observed relatively small vascular networks at day 14, that grew increasingly complex through days 21 and day 28 (Fig. 1E–G). Co-staining with LIM homeobox 1 (LHX1; nephron progenitor cells) and ECAD indicated preserved epithelial patterning alongside vascular outgrowth.

### Quantitative 3D Image Analysis Confirms Increased Vessel Complexity in HC-Treated Organoids

We next quantified the impact of HC treatment on vascular development, using IMARIS software to assess the increase in percent vessel volume (relative to total organoid volume), total vessel length, and the number of branch points compared to CON (Fig. 2A). HC-A-treated organoids exhibited significantly increased vessel volume on days 14, 21, and 28 (Fig. 2B), accompanied by longer vessel length and greater branch complexity on days 21 and 28 (Fig. 2C, D).

**Figure 2:**
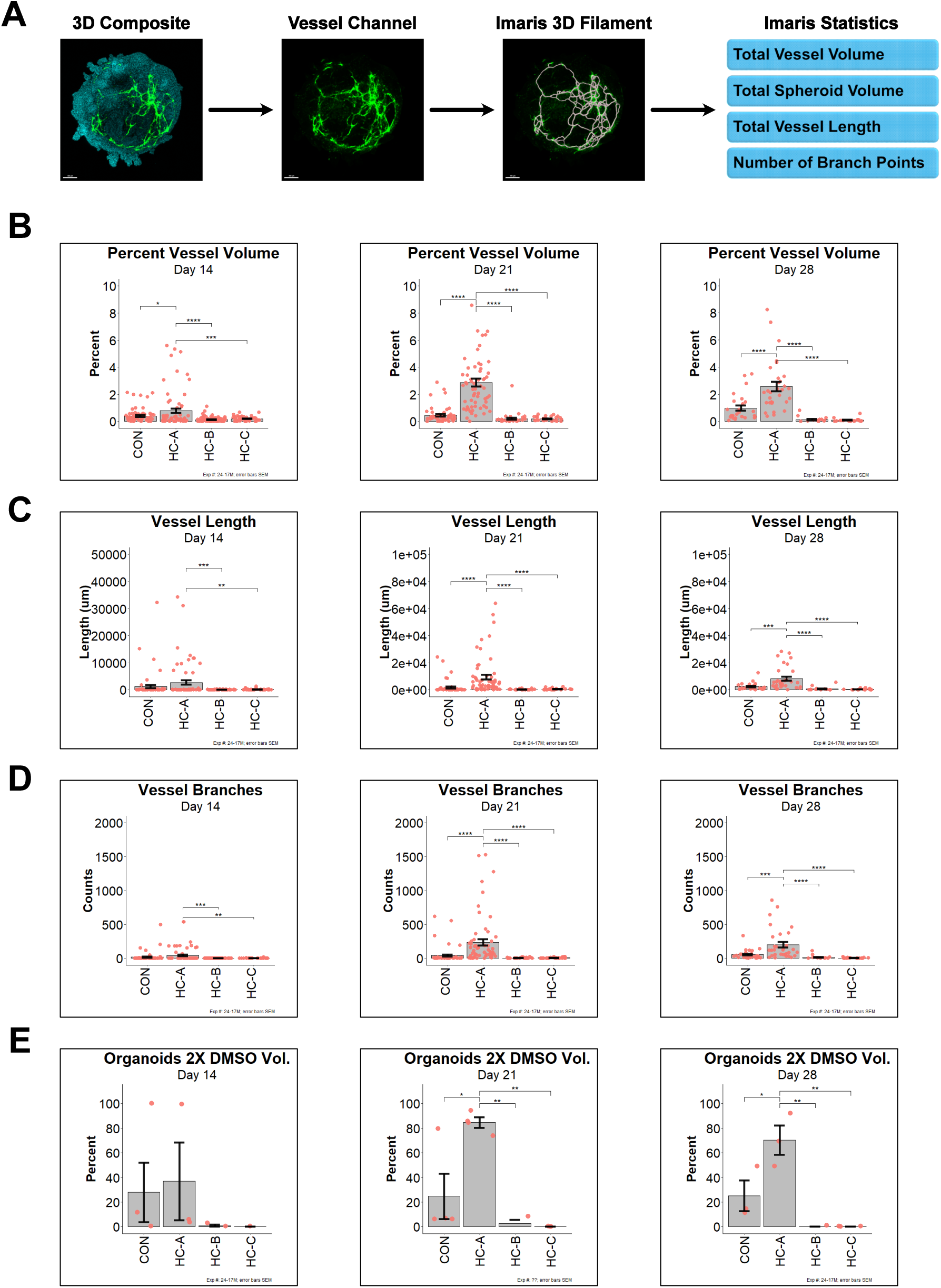
Three-dimensional volumetric analysis of HC-induced vessels in kidney organoids. (A) Image analysis workflow in IMARIS. DAPI channel was used for total spheroid volume calculation and the CD31 channel was isolated for vascular filament analysis. (B) Quantification of vessel volume as a percentage of the total organoid volume, per organoid on day 14 (left panel), day 21 (middle panel), and day 28 (right panel). (C) Quantification of the total vessel length per organoid on day 14 (left panel), day 21 (middle panel), and day 28 (right panel). (D) Quantification of the total number of vessel branches per organoid on day 14 (left panel), day 21 (middle panel), and day 28 (right panel). (E) Quantification of the percentage of organoids with at least twice the mean percent vessel volume found in control organoids on day 14 (left panel; *n=3* biological replicates), day 21 (middle panel; *n=3* biological replicates), and day 28 (right panel; *n=3* biological replicates). Day 14 sample size: CON *n=65* organoids combined across 3 replicates, HC-A *n=67* organoids combined across 3 biological replicates, HC-B *n=80* organoids combined across 3 biological replicates, and HC-C *n=48* organoids combined across 3 biological replicates. Day 21 sample size: CON *n=53* organoids combined across 3 biological replicates, HC-A *n=61* organoids combined across 3 biological replicates, HC-B *n=32* organoids combined across 3 biological replicates, and HC-C *n=32* organoids combined across 3 biological replicates. Day 28 sample size: CON *n=26* organoids combined across 3 biological replicates, HC-A *n=31* organoids combined across 3 biological replicates, HC-B *n=17* organoids combined across 3 biological replicates, and HC-C *n=20* organoids across 3 biological replicates. Data are represented as mean ± SEM. * p< 0.05, ** p < 0.01, *** p < 0.001, and **** p < 0.0001.

We also assessed the robustness and variability of HC’s effect on vascular development. We analyzed the percentage of organoids that exhibited a percent vessel volume of at least 2-fold higher than the average percent vessel volume found in our controls (Figure 2E). Here we also observed that HC-A-treated organoids had a significantly higher percentage of highly vascularized organoids. For each of these measurements of vascular development, we saw no significant increases in HC-B-treated or HC-C-treated organoids compared to controls at any timepoint. From this point forward, we characterized HC-induced EC differentiation using only the HC-A protocol (HC hereafter).

### Both GR and MR Contribute to Hydrocortisone-Induced Endothelial Differentiation

We next investigated the receptor-specific contributions of HC, by combining HC treatment with either a GR or an MR inhibitor (GRi or MRi). We found that organoids co-treated with HC and GRi exhibited reduced CD31 expression across all timepoints, with the effects becoming more pronounced with time (Fig. 3A, B), suggesting that the response is mediated at least in part by GR. Surprisingly, we also observed that organoids co-treated with HC and MRi also reduced CD31 protein levels at all time points, although these results are less striking (Fig. 3C, D).

**Figure 3:**
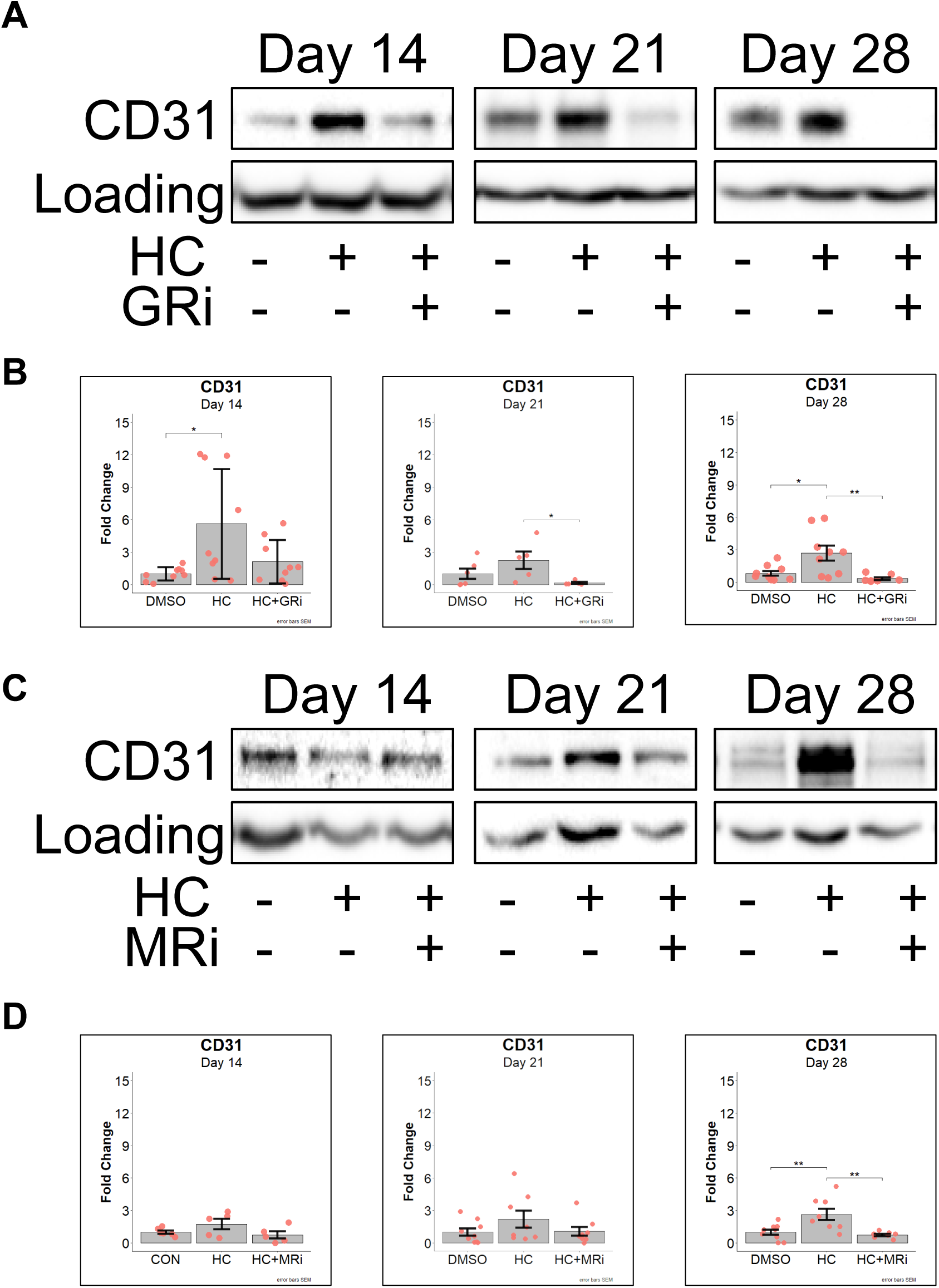
Analysis of GR and MR contribution to EC differentiation in kidney organoids. (A) Representative western blots of CD31 and GAPDH (Loading) protein expression at day 14 (left panels), day 21 (middle panels), and 28 (right panels). (B) Densitomitry quantification of CD31 protein expression from panel A. (C) Representative western blots of CD31 and GAPDH (Loading) protein expression at day 14 (left panels), day 21 (middle panels), and 28 (right panels). (D) Densitomitry quantification of CD31 protein expression from panel C. *n=9* biological replicates combined across 3 experiments per treatment group (B and D). Data are represented as mean ± SEM. * p< 0.05, ** p < 0.01, *** p < 0.001, and **** p < 0.0001.

To determine the impact of receptor blockade on vessel formation, we employed IMARIS to analyze vessel morphology as described above. We found that by day 14 both GR and MR inhibition significantly reduced percent vessel volume, total vessel length, and number of branch points (Supplemental Fig. 2A, B). However, because our HC-treated and control organoids have similar percent vessel volume by day 14 of differentiation, we did not see a reduction in the percentage of highly vascularized organoids under either GR or MR inhibition (Supplemental Fig. 2A, B). Together, these findings suggest that both GR and MR receptors are involved in mediating HC-induced EC differentiation.

### Pathway Analysis Highlights Early Transcriptional Reprogramming by HC

To elucidate transcriptional changes underlying HC-induced EC differentiation, we performed bulk RNA-seq at day 9—the final day of HC treatment. Gene set enrichment analysis (GSEA) of the day 9 RNA-seq dataset revealed significant upregulation of pathways associated with MYC targets, oxidative phosphorylation, and downregulation of inflammation (Supplemental Fig. 3A), all of which were expected after treatment with HC. Surprisingly, we found that the angiogenesis pathway was also downregulated (Supplemental Fig. 3A). We then performed TISSUES ontology analysis to identify the cell and tissue types predicted to be downregulated (Palasca et al., 2018). We found that EC-related tissues comprised six out of the top 10 negatively enriched tissues (Supplemental Fig. 3B). These data indicate that HC suppresses early transcriptional programming of many canonical EC signatures.

### Transcriptomic Profiling Reveals Inhibition of Angiogenic Gene Programs by Hydrocortisone

A heatmap of the top 50 differentially expressed genes enriched for the TISSUES:Endothelium ontology highlighted broad downregulation of known angiogenic genes following HC treatment (Fig. 4A). Canonical EC junction genes such as *vecad* and *mcam* were among the most significantly reduced, while EC differentiation factor genes, such as *ang1* and *vegfa*, were slightly upregulated (Fig. 4B). Validation via RT-qPCR confirmed *ang1* mRNA levels were increased following HC treatment but revealed that *vegfa* gene expression was downregulated (Fig. 4C).

**Figure 4:**
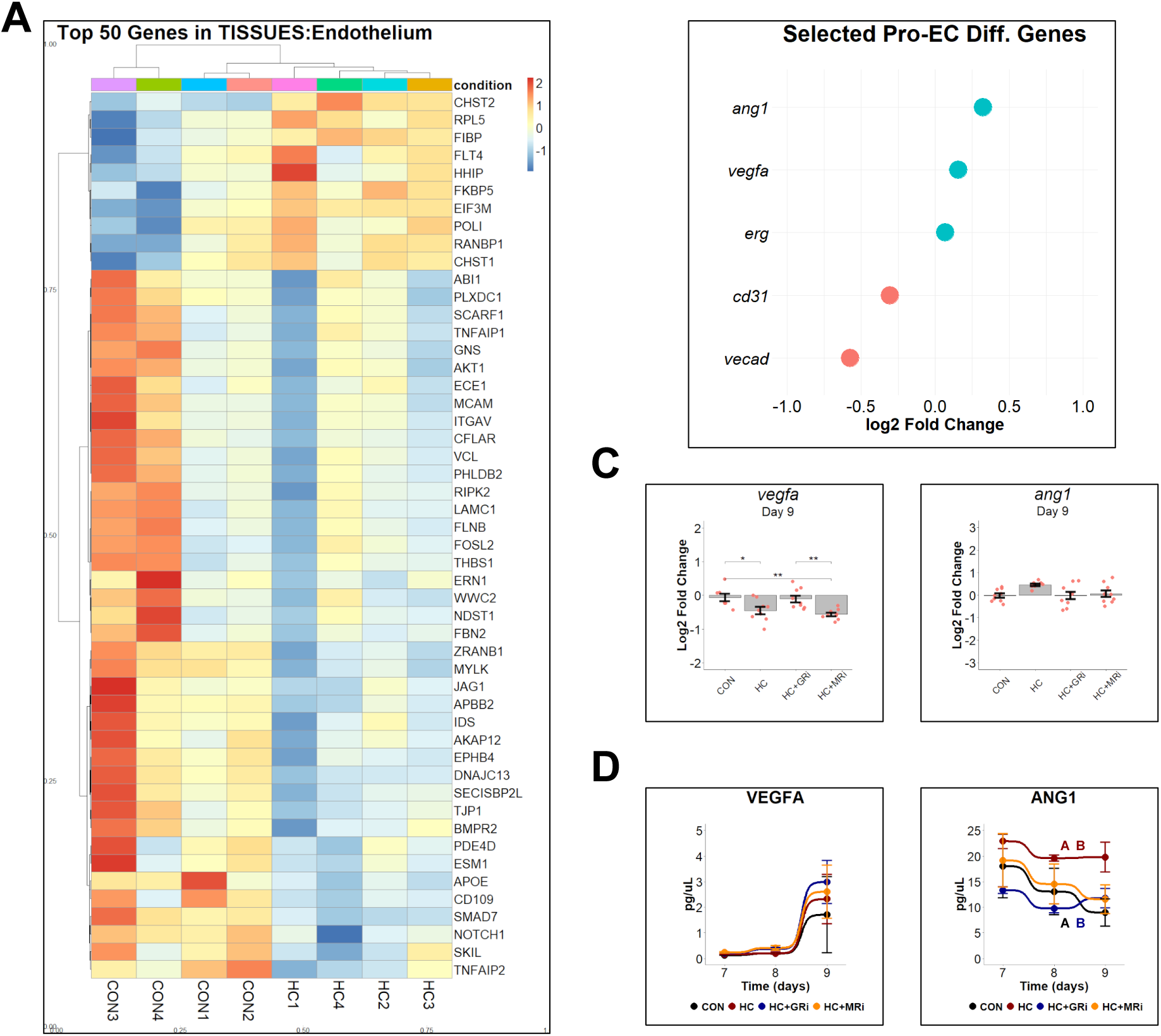
Identification and analysis of EC differentiation factors. (A) Heatmap of the top 50 DEGs found overlapping the TISSUES:Endothelium ontology. (B) Dotplot of the Log2 fold change (DMSO- vs HC-treated) in mRNA expression of selected EC genes from bulk RNA-seq on day 9. (C) Log2 fold change in mRNA expression on day 9 by RT-qPCR for *vegfa* (left panel) and *ang1* (right panel). (D) Quantification of secreted VEGFA protein (left panel) and ANG1 (right panel) via ELISA from cell culture supernatants on days 7, 8, and 9 of differentiation. *n=9* biological replicates combined across 3 experiments per treatment group (C-D). Data are represented as mean ± SEM. * p< 0.05, ** p < 0.01, *** p < 0.001, and **** p < 0.0001. ^A^ p<0.05, ^B^ p<0.05; color represents compared groups.

We then asked if the levels of *vegfa* and *ang1* secreted proteins reflect their gene expression patterns. Indeed, we found enhanced secretion of ANG1 protein on days 7–9, but no change in VEGFA protein secretion during that time (Fig. 4D). These results suggest that HC drives upregulation of pro-EC survival factors but may inhibit vessel formation and branching during early time points.

### HC Treatment Preserves Nephrogenic Program and Limits Pro-Inflammatory Gene Expression

Assessment of gene expression on day 24 confirmed maintenance of both vascular and nephron lineages following HC treatment. Specifically, EC junction markers (*cd31* and *vecad*), a hematopoietic marker (*cd34*), and renal markers including *aqp2*, *gata3*, *pax2*, *wt1*, and *ren* remained detectable following HC treatment (Supplemental Fig. 1A, C). The inflammatory cytokines *tnfα*, *il6*, and *ccl2* were left unchanged (Supplemental Fig. 1D), suggesting no impact on inflammatory activation.

Gene expression analysis at day 9—the end of HC treatment—further confirmed preserved nephrogenic potential following HC treatment. We found, through bulk RNA-seq, that selected mesoderm progenitor and nephrogenic genes either remained stable or were upregulated in HC-treated cultures (Supplemental Fig. 4A). However, these results were not recapitulated by RT-qPCR. Our RT-qPCR results showed that while *osr1* expression was unchanged, *lhx1*, *six2*, and *pax2* tended to decrease on average, although these results were variable and not statistically significant (Supplemental Fig. 4B). Finally, we observed via RT-qPCR that HC treatment (alone or in combination with GRi or MRi) resulted in a small, statistically insignificant decrease in *tnfα* and *il6* gene expression (Supplemental Fig. 4C). Together, these data indicate that HC promotes vascular induction without completely compromising nephron identity.

## Discussion

Our findings reveal a previously underappreciated role for glucocorticoid signaling in orchestrating vascular development within human kidney organoids. Specifically, we demonstrate that hydrocortisone (HC), a synthetic cortisol analog, enhances endothelial cell (EC) differentiation and promotes complex vessel formation when administered during early kidney organoid patterning. These effects are robust, reproducible, and observed across multiple morphological and molecular endpoints including increased vascular volume and branch complexity, and upregulation of key endothelial markers. Importantly, we show that these effects occur without disrupting nephron identity, positioning glucocorticoid signaling as a potent and tractable modulator of vascular development *in vitro*.

A particularly compelling finding of our study is that HC-induced vascular development is mediated by the cooperative activity of both the glucocorticoid receptor (GR) and the mineralocorticoid receptor (MR). While GR has been broadly implicated in endothelial biology, including EC homeostasis (Przybyciński et al., 2021; Srivastava et al., 2021), permeability (Salvador et al., 2014), and response to stress (Zielińska et al., 2016), its role in vascular development remains less clearly defined. MR, traditionally associated with ion homeostasis, has recently emerged as a context-dependent regulator of EC behavior (Barrera-Chimal and Jaisser, 2021; Camarda et al., 2025; Gorini et al., 2019; Ibarrola and Jaffe, 2024; Lother et al., 2019), yet few studies have explored its developmental functions (McCann et al., 2021; Young and Clyne, 2021). Our observation that inhibition of either GR or MR impairs HC-induced EC differentiation underscores the necessity of GR-MR crosstalk in coordinating pro-vascular outcomes. Recent research suggests that GR and MR cooperate to regulate downstream transcription (Johnson et al., 2024). Together, these data support a model in which glucocorticoids act through both receptors to fine-tune transcriptional programs essential for endothelial specification and morphogenesis.

Interestingly, our protein-level analyses revealed that HC-treated organoids exhibited robust vascular development by day 21, even though HC treatment suppressed canonical angiogenic gene programs on day 9. Our results suggest that HC treatment appears to initiate a distinct EC differentiation trajectory characterized by downregulation of junctional genes such as *vecad* and *mcam*, while modestly upregulating pro-survival and stabilization factors like *ang1*. These findings suggest that HC does not simply accelerate angiogenesis via conventional VEGFA-dependent pathways but instead primes endothelial progenitors by establishing a transcriptional state conducive to long-term vessel stabilization and complexity. The increased secretion of ANG1, in the absence of elevated VEGFA, further supports a model in which HC favors endothelial survival and quiescence over sprouting angiogenesis during early stages of differentiation.

While our results highlight the vascular potential of HC, they also underscore its selectivity. Importantly, we observed preservation of nephrogenic markers in HC-treated organoids, including podocyte and tubular epithelial genes (Supplemental Fig. 1 and 4), throughout the differentiation process. This is a key consideration in the development of vascularized kidney organoids, as excessive activation of EC pathways often compromises epithelial lineage fidelity (Perens et al., 2016). Additionally, we found no evidence—at the transcriptional level—for broad inflammatory activation following HC exposure, alleviating concerns about off-target stress responses.

Taken together, our findings establish glucocorticoids as powerful regulators of vascular development in kidney organoids and open new avenues for exploring hormone-driven control of cell fate specification. However, the precise molecular mechanisms by which GR and MR orchestrate EC differentiation remain to be elucidated. While there is evidence that GR and MR can enhance transcriptional output of pro-EC transcription factors (Srivastava *et al*., 2019), it will be critical to define the direct transcriptional targets of each receptor and the potential for receptor co-occupancy or competition on chromatin. Furthermore, the temporal windows within which GR and MR exert distinct or overlapping functions warrant further exploration.

In conclusion, this study identifies glucocorticoid signaling as a potent modulator of endothelial differentiation in human kidney organoids. By revealing an unexpected developmental role for GR and MR in vascular patterning, our work provides a foundation for future studies aimed at mechanistically dissecting steroid hormone signaling in organogenesis. These insights have broad implications not only for the engineering of vascularized tissues but also for understanding how hormonal cues shape the development of the human vasculature.

### Limitations of the Study

The mechanism by which glucocorticoid signaling contributes to EC differentiation and vascular development described in this work is incomplete. We characterized the contribution of HC receptors (GR and MR) to EC differentiation and vessel formation, but we did not directly address whether HC-induced expression of ANG1 directly contributes to vascular development. While glucocorticoid signaling is important for EC resilience *in vivo* (Srivastava *et al*., 2021), its role(s) in EC differentiation *in vivo* remain a mystery. Finally, it is important to note that excess maternal dexamethasone exposure—overactivation of the GR—promotes adverse developmental defects in the embryonic kidney (Wang et al., 2022). Therefore, while we assessed how HC treatment modulated expected gene expression patterns for kidney-related genes, we did not deeply characterize these changes or assess functionality following HC treatment. Moving forward, we envision that vascularized kidney organoids will provide a powerful *in vitro* platform to explore the interactions between multiple tissues during development in humans. By combining human organoid studies with animal models, we will be able to identify important conserved signals that could be exploited to ameliorate disease.

## STAR Methods

### Key Resources Table

| REAGENT or RESOURCE | SOURCE | IDENTIFIER |
| --- | --- | --- |
| **Antibodies** |  |  |
| Sheep Anti-CD31 (PECAM1) | R&D Systems | AF806 |
| Goat Anti-VE-Cadherin | R&D Systems | AF938 |
| Mouse Anti-LHX1 | DSHB | 4F2-C |
| Mouse Anti-E-Cadherin | BD Biosciences | 610182 |
| Rabbit Anti-GAPDH | Abcam | AB181602 |
| Goat Anti-Rabbit IgG (H+L) Secondary Antibody, HRP | Invitrogen | 65-6120 |
| Donkey Anti-Sheep IgG (H+L) Secondary Antibody, HRP | Invitrogen | A16041 |
| Rabbit Anti-Goat HRP Conjugated Antibody | R&D Systems | HAF017 |
| **Bacterial and virus strains** |  |  |
| **Biological samples** |  |  |
| **Chemicals, peptides, and recombinant proteins** |  |  |
| Geltrex™ LDEV-Free Reduced Growth Factor Basement Membrane Matrix | Gibco | A1413202 |
| StemFlex™ Medium | Gibco | A3349401 |
| DPBS, no calcium, no magnesium | Gibco | 14190144 |
| StemPro™ Accutase™ Cell Dissociation Reagent | Gibco | A1110501 |
| DMEM/F-12, no glutamine | Gibco | 21331020 |
| Y-27632 dihydrochloride | R&D Systems | 1254/10 |
| Advanced RPMI 1640 Medium | Gibco | 12633020 |
| L-Glutamine (200 mM) | Gibco | 25030081 |
| KnockOut™ Serum Replacement | Gibco | 10828028 |
| CHIR 99021 | Tocris | 4423 |
| Human Noggin Recombinant Protein | Peprtech | 120-10C |
| Recombinant Human/Mouse/Rat Activin A Protein | R&D Systems | 338-AC-500/CF |
| Heparin sodium salt from porcine intestinal mucosa | Sigma-Aldrich | H3149 |
| Poly(vinyl alcohol) | Sigma-Aldrich | 363081 |
| Methyl cellulose | Sigma-Aldrich | M7027 |
| Recombinant Human FGF-9 Protein | R&D Systems | 273-F9-025 |
| Hydrocortisone | Tocris | 4093 |
| Mifepristone | Tocris | 1479 |
| RU28318 | Tocris | 76676-34-1 |
| cOmplete™ Protease Inhibitor Cocktail | Sigma-Aldrich | 11697498001 |
| PhosSTOP™ | Sigma-Aldrich | 4906845001 |
| EveryBlot Blocking Buffer | Bio Rad | 12010020 |
| Precision Plus Protein™ All Blue Prestained Protein Standards | Bio Rad | 1610373 |
| Clarity Western ECL Substrate | Bio Rad | 1705061 |
| Western Blot Stripping Buffer | Abcam | AB282569 |
| Donkey Serum | Sigma-Aldrich | D9663 |
| Formamide | Ambion | AM9342 |
| Paraformaldehyde 16% Aqueous Solution EM Grade | EMS | 15710 |
| Penicillin (10,000 units)-Streptomycin (10mg) | Sigma-Aldrich | P4333 |
| **Critical commercial assays** |  |  |
| Roche LightCycler 480 Probes Master Mix | Roche | 04707494001 |
| RNeasy Mini Kit | Qiagen | 74106 |
| SuperScript IV First-Strand Synthesis System | Invitrogen | 18091200 |
| NEBNext® Poly(A) Magnetic Isolation Module | NEB | E7490 |
| NEBNext® Ultra II RNA Library Prep Kit for Illumina® | NEB | E7770S |
| NEBNext® Multiplex Oligos for Illumina® (Index Primers Set 1) | NEB | E7335S |
| Pierce™ BCA Protein Assay Kits | ThermoFisher | A65453 |
| **Deposited data** |  |  |
| **Experimental models: Cell lines** |  |  |
| Gibco™ episomal hiPSC line | Gibco | A18945 |
| **Experimental models: Organisms/strains** |  |  |
| **Oligonucleotides** |  |  |
| TaqMan™ Gene Expression Assays GAPDH | ThermoFisher | Hs02786624_g1 |
| TaqMan™ Gene Expression Assays CDH1 (ECAD) | ThermoFisher | Hs01023895_m1 |
| TaqMan™ Gene Expression Assays GATA3 | ThermoFisher | Hs00231122_m1 |
| TaqMan™ Gene Expression Assays PAX2 | ThermoFisher | Hs01057416_m1 |
| TaqMan™ Gene Expression Assays NPHS1 | ThermoFisher | Hs00190446_m1 |
| TaqMan™ Gene Expression Assays WT1 | ThermoFisher | Hs01103751_m1 |
| TaqMan™ Gene Expression Assays AQP2 | ThermoFisher | Hs00166640_m1 |
| TaqMan™ Gene Expression Assays REN | ThermoFisher | Hs00982555_m1 |
| TaqMan™ Gene Expression Assays CD34 | ThermoFisher | Hs02576480_m1 |
| TaqMan™ Gene Expression Assays TNF | ThermoFisher | Hs00174128_m1 |
| TaqMan™ Gene Expression Assays IL6 | ThermoFisher | Hs00985639_m1 |
| TaqMan™ Gene Expression Assays CDH5 (VECAD) | ThermoFisher | Hs00901465_m1 |
| TaqMan™ Gene Expression Assays PECAM1 (CD31) | ThermoFisher | Hs01065279_m1 |
| TaqMan™ Gene Expression Assays CCL2 | ThermoFisher | Hs00234140_m1 |
| TaqMan™ Gene Expression Assays VEGFA | ThermoFisher | Hs00900055_m1 |
| TaqMan™ Gene Expression Assays OSR1 | ThermoFisher | Hs01586544_m1 |
| TaqMan™ Gene Expression Assays SIX2 | ThermoFisher | Hs00232731_m1 |
| TaqMan™ Gene Expression Assays LHX1 | ThermoFisher | Hs00232144_m1 |
| TaqMan™ Gene Expression Assays ANGPT1 (ANG1) | ThermoFisher | Hs00919202_m1 |
| **Recombinant DNA** |  |  |
| **Software and algorithms** |  |  |
| Fiji | Schindelin, J. (PMID: 22743772) | https://imagej.net/software/fiji/ |
| IMARIS | Oxford Instruments | V10.2 |
| R | R Core Team (2023) | https://www.R-project.org/ |
| R Studio | Posit team (2023) | http://www.posit.co/ |
| **Other** |  |  |

## Resource Availability

### Lead Contact

Further information and requests for resources and reagents should be directed to and will be fulfilled by the lead contact, Cory P. Johnson.

### Materials Availability

No unique materials/reagents have been generated in this study.

### Data and Code Availability

The bulk RNA sequencing data generated in this study have been submitted to NCBI Gene Expression Omnibus (GSE304936). All code used to generate and analyzed data presented in this article are available upon request.

### Human Pluripotent Stem Cell Culture and Maintenance

All experiments were performed using Gibco episomal iPS cells (Gibco, # A18945) cultured per manufacturer’s instructions. Cells were seeded into experimental plates at 20,000-40,000 cells/cm^2^ in StemFlex (Gibco, # A3349401) containing 10uM Y-27632 dihydrochloride (R&D, #1254) and 1% penicillin-streptomycin (Sigma, #P4333) and seeded on plates coated with 1% Geltrex (Gibco, # A1413202). All cell cultures were maintained at 37°C and supplemented with 5% CO_2_.

### Renal Organoid Differentiation and Culture

Organoids were differentiated using a modified combination of protocols from Morizane et al. and Kumar et al (PMID: 26458176, PMID: 30846463). Day 0 cells, plated on Day -1 at ∼40,000 cells/cm2, were washed with pre-warmed 1X dPBS and switched to differentiation media Advanced RPMI 1640 (Gibco, #12633020) + 1% L-Glutamine (Gibco, #25030081) + 0.5% KnockOut Serum Replacement (Gibco, #10828028), containing 7-10μM CHIR (Tocris, #4423) and 5ng/mL noggin (Peprotech, #120-10C) on days 0-3 with media changes every 2 days. On day 4, cells received differentiation media containing 10ng/mL Activin A (R&D, #338-AC-500/CF) and 1μg/mL heparin (Sigma, #H3149). At day 7, media was replaced with differentiation media containing 10ng/mL FGF9 (R&D, #273-F9-025) and 1μg/mL heparin. On day 9, cells received media containing 10ng/mL FGF9 (R&D, #273-F9-025), 3μM CHIR, and 1μg/mL heparin. On day 11, cells were dissociated into single-cell suspensions using Accutase, pelleted at 300g for 3 minutes, then resuspended in differentiation medium containing 0.1% polyvinyl alcohol (PVA), 0.1% methylcellulose (MC), 1μg/mL heparin, and 10μM Y-27632 (Tocris, #1254) equivalent to a splitting ratio of 1:2-1:4. Cells were then transferred to cell repellent dishes (Grenier Bio-One, #657970) and placed on an orbital shaker (Benchmark, #BT40001) at 60-80rpm in a cell culture incubator overnight to form spheroids. Day 12 spheroids were collected in 15mL conical tubes, centrifuged at 10g for 1 minute, and treated with fresh differentiation medium containing 0.1% PVA and 0.1% MC, without Y-27632 or growth factors. Subsequent media changes were conducted every other day until collection.

### Hydocortisone and other Drug Treatments

Hydrocortisone (HC) was used at a final concentration of 1ug/mL for all HC treatments. All daily drug treatments were conducted with media changes each day containing the prescribed growth factors appropriate to each time point. Glucocorticoid receptor inhibitor RU486 (Tocris; CAT#: 1479) and mineralocorticoid receptor inhibitor RU28318 (Tocris; CAT#: 76676-34-1) were used at a final concentration of 10uM.

### Total RNA Isolation and cDNA Synthesis

Total RNA was isolated per manufacturer’s instructions using the RNeasy RNA Isolation Kit (Qiagen, #74104). RNA quantity and purity were analyzed using the “Nanodrop”. For cDNA synthesis, RNA input was normalized across all samples. Synthesis of cDNA was conducted using the SuperScript IV First-Strand Synthesis System (Invitrogen, #18091200) per manufacturer’s protocol.

### RT-qPCR

All gene expression assays were conducted using pre-designed TaqMan probes from ThermoFisher. For each sample, 20ng/uL of cDNA template was added per gene and all target genes were normalized to a GAPDH housekeeper gene. All gene expression assays were run using a Roche LightCycler 480 in Roche LightCycler 480 Probes Master Mix (Roche; CAT#: 04707494001) according to manufacturer’s instructions. Cp values were calculated using the XX built-in software’s “2^nd^ Derivative/Abs Quant” function followed by conversion to 2^-ΔΔCp^.

### Bulk RNA Sequencing Library Preparation and Sequencing

Organoids were differentiated in monolayer culture to day 9 and RNA was isolated per manufacturer’s instructions using the RNeasy RNA Isolation Kit (Qiagen, #74104). Isolation of Poly(A) mRNA was conducted using the NEBNext Poly(A) Magnetic Isolation Module (NEB, #E7490). Following isolation of poly(a) mRNA, library preparation was conducted using the NEBNext Ultra II RNA Library Prep Kit for Illumina (NEB, #E7770S) per manufacturer’s instructions, individual samples were barcoded using NEBNext Multiplex Oligos for Illumina (NEB, #E7335S), and equal sample volumes were pooled prior to sample QC. Quantities of the final libraries were assessed by Qubit 2.0 (ThermoFisher) and QuantStudio ® 5 System (Applied Biosystems) and quality was assessed by TapeStation D1000 ScreenTape (Agilent Technologies Inc.). Libraries were loaded based on QC values and sequenced on an Illumina® *NovaSeq X Plus 10B* (Illumina) with a read length configuration of *150 PE* for a total of 2-2.5 B PE reads (*1-1.25*B reads in each direction).

### Bulk RNA Sequencing Analysis

Sequence files were analyzed with the Nextflow (https://nextflow.io/) NF-core (https://nf-co.re/) rnaseq workflow, version 3.18.0 (https://nf-co.re/rnaseq/3.8.0), using the Memverge Opcenter interface to Amazon Web Services computing. Version of all software included in the rnaseq v3.18.0 are listed in supplemental file nf_core_rnaseq_software_mqc_versions.yml and all parameters of the analysis in the supplemental file params_2025-03-07_17-39-07.json. Sequences were aligned to the human genome build GRC38 with annotations drawn from Ensembl release 113 (Dyer et al., 2024). The count matrix was filtered for genes with a summed expression total greater than 20 across the four replicates of at least one condition. Differential expression analysis was carried out with R package DESeq2 version 1.44.0 (Love et al., 2014), controlling for any systematic replicate effect with the design equation ‘∼replicate + treatment.’

### Functional Annotation and Pathway Analysis

Gene interaction networks and tissue enrichment data were obtained using STRINGdb to query the TISSUES database (Szklarczyk et al., 2023). This analysis provided predicted enrichment of specific tissue types using our entire gene expression dataset ranked by log2 fold change values for each gene. These data were subsequently visualized using heatmaps generated with the pheatmap package and additional plots created with ggplot2 (Kolde, 2019; Wickham et al., 2016).

Gene ontology (GO) and pathway enrichment analyses were performed using the msigdbr package to access gene sets from the Molecular Signatures Database (MSigDB). Gene Set Enrichment Analysis (GSEA) was conducted using the fgsea package in R, with genes ranked by log2 fold change (Castanza et al., 2023; Liberzon et al., 2015; Subramanian et al., 2005). Enrichment significance was determined using an adjusted p-value < 0.05 and a normalized enrichment score (NES) of ± 0.5.

### Immunostaining, Clearing, and Mounting

Organoids were fixed in 4% PFA for 1 hour at room temperature on a microtube rotator. Following fixation, organoids were washed 3 times with 1X PBS + 0.01% Triton X-100 for 20 minutes at room temperature. Next, organoids were incubated in blocking solution (5% BSA, 5% Donkey Serum, 0.1% Triton X-100 in PBS) overnight, but no more than 3 days, at 4°C on a microtube rotator. After blocking, organoids were incubated with primary antibodies (1:100 dilution) in fresh blocking solution for up to 2 days at 4°C on a microtube rotator. Following primary antibody incubation, organoids were washed 3 times for 20 minutes each at room temperature in PBS + 0.01% Triton X-100 (PBST). Next, organoids were incubated with secondary antibodies in blocking solution overnight at 4°C on a microtube rotator. Following secondary incubation, organoids were washed 2 times for 20 minutes each at room temperature with PBST and a third wash in PBS. Organoids were then stained with DAPI at 2ug/mL in PBS for 7-10 minutes at room temperature and washed 3 times with 1X PBS. Immunostained organoids were cleared using a formamide gradient and mounted using Aquapolymount (Polysciences, #18606-20). The formamide gradient was as follows: 30 minutes in 20% formamide, 30 minutes in 40% formamide, 1 hour in 80% formamide, 30 minutes in 95% formamide, and then indefinitely in 95% formamide.

### Volumetric Image Acquisition

Images were acquired using a Spinning-disk confocal unit (Yokogawa, #CSU-W1) on a Nikon inverted Ti-Eclipse microscope stand (Nikon, #TI-E-736803), equipped with a CFI Plan Apochromat Lambda D 10x/0.45 objective lens (Nikon, #MRD70040).

DAPI, Alexa 488, Alexa 594, and Alexa 647 fluorescence were excited with 405nm, 488nm, 561nm, and 640nm respectively. All lasers were used at 100% from a LUNF-XL laser combiner (Nikon, #77098033) and collected using a DM 405/488/561/640 dichroic beamsplitter (Nikon, #MHE46420) with ET436/20x single-band bandpass (Chroma, #ET436/20x), 525/50 nm BrightLine® single-band bandpass (Semrock, #FF01-525/50), ET605/52m bandpass (Chroma, #ET605/52m), and ET705/72m single-band bandpass (Chroma, #ET705/72m) emission filters respectively.

Images were acquired in 2048*2048 pixels, at 10x magnification with a CMOS Zyla 4.2 camera (Andor) controlled with NIS AR 5.41.02 (Nikon, build 1711) software and saved in Nd2 file format.

### Analysis of Percent Vascular Volume, Branching, and Length

Spheroid images were converted to .IMS file format using the IMARIS file converter software. In IMARIS, DAPI-stained nuclei were used to generate a 3D volumetric surface rendering to quantify total spheroid volume. Images were batch processed using a 5um gaussian filter, surface grain size of 1.28um, and manual threshold value of 105. After surface creation, objects less than 2.00e-7 um^3 were filtered. Next CD31+ vessels were used to generate 3D volumetric filaments to quantify vascular volume. Images were batch processed using a 3μm gaussian filter. Filaments were created using the Autopath Algorithm to detect filaments with loops but no somas or spines. Seed points were set to 1.92um for thinnest diameter and 100μm for largest diameter. Chosen seed points were a minimum pixel value of 100 and segment diameter smoothing was set to medium. Finally, filaments were segmented on a trained segment classification model using a maximum gap length of 2μm between segments. The resulting cumulative vascular volume was then divided by total spheroid volume and multiplied by 100 to determine the percent vascular volume for each spheroid.

### Western Blotting

Cell lysis and protein extraction were conducted as follows: Spheroids were centrifuged at 16,000g for 15 seconds and supernatant was removed. Organoids were suspended in lysis buffer (50mM Tris pH 8, 150mM NaCl, 0.5% SDS, and 0.5% Sodium Deoxycholate) containing protease (SIGMA, #11697498001) and phosphatase inhibitors (SIGMA, #4906845001) and frozen at -20°C overnight. Following freezing, organoid lysate was thawed on ice, further homogenized with a pestle, and then centrifuged at 4°C and 16,000g for 20 minutes to remove cell debris. The supernatant was transferred to a new microcentrifuge tube and protein concentration was measured using the Pierce BCA Protein Assay Kit (ThermoFisher, #A65453). Protein samples were normalized to a minimum of 5ug total protein per sample volume in sample buffer. Samples were then boiled at 95°C for 5 minutes and either used immediately or stored at -20°C for long-term storage. SDS-PAGE gels were made to a 10% final concentration of acrylamide. Gels were run using protein running buffer (2.9g Tris base, 14.4g Glycine, and 1g SDS per liter) at 60V for 20 minutes followed by 100V until the dye front reached the bottom of the gel. Protein was transferred using standard wet transfer (BioRad, #1658033) for 90 minutes at 100V to a PVDF membrane. Following transfer, membranes were stained for total protein with Ponceau S stain for 5 minutes, destained in 10% acetic acid for 5 minutes, and imaged using an iPhone camera. Membranes were blocked using EveryBlot Blocking Buffer (BioRad, #12010020) for 20 minutes at room temperature. Primary antibodies were diluted in blocking buffer and membranes were incubated at 4°C overnight while rocking. Protein size confirmation was identified using the Precision Plus All Blue Pre-stained Protein Ladder (BioRad, #1610373). Membranes were incubated with the respective secondary antibodies on a rocker at room temperature for 90 minutes followed by two 10-minute washes in PBS + 0.1% Tween 20 and one 10-minute wash in PBS. All secondary antibodies were used at 1:3000 dilution. Finally, all membranes were incubated with ECL substrate (BioRad, #1705061) for a minimum of 1 minute per manufacturer’s instructions and imaged using the Syngene G:Box Chemi-XRQ (Syngene) imaging machine. Stripping and reprobing was conducted per manufacturer’s instructions (Abcam, #AB282569).

## Acknowledgements

The authors wish to thank Yullia Kiian, Ph.D. and Ekaterina Pecksen, Ph.D., for their training in organoid culture; Patricia Schroeder, Ph.D., for helping with sample collection; Iain Drummond, Ph.D., James Coffman, Ph.D., James Godwin, Ph.D., Caramai Kamei, Ph.D., Anastasia Paulmann, M.D., Friedrich Luft, M.D., Ron Korstanje, Ph.D., Prayag Murawala, Ph.D., and William Sampson for critical feedback and discussion; Marko Pende, Ph.D., for critical feedback and discussion regarding organoid clearing; Frederic Bonnet, Ph.D. and the Light Microscopy Facility (https://lmf.mdibl.org/) for training, critical discussion and feedback; and Stephen B. Sampson for manuscript editorial feedback and comments. Research reported in this publication was supported by an Institutional Development Award (IDeA) from the National Institute of General Medical Sciences of the National Institutes of Health under grant numbers P30GM154610, P20GM203423, 1P20GM144265-01A1, and the Scott R. Mackenzie foundation. Additional support from NSF under grant numbers 2243416 and OIA-1849227, and the MDI Biological Laboratory.

## Author Contributions

CPJ wrote/edited the original manuscript and designed, conducted, analyzed, and oversaw all experiments. HMS edited the manuscript and conducted and analyzed experiments. HF and JG conducted DEG analysis. LBS conducted experiments and collected samples. KET conducted experiments and analyzed data. SEC and SB conducted experiments and collected samples. CHT conducted experiments. MDC reviewed the manuscript, discussed bulk RNA sequencing functional analysis approaches, and assisted with data submission. MM reviewed and edited the manuscript. HH edited the manuscript and oversaw the project.

## Declaration of generative AI and AI-assisted technologies

ChatGPT (OpenAI, v4 and v4.5) and Claude AI (Anthropic, Sonnet 4) were used to edit and refine the manuscript for clarity and word limit. All AI-based edits were extensively reviewed and further edited for accuracy and clarity.

---

**Supplemental Figure 1:**
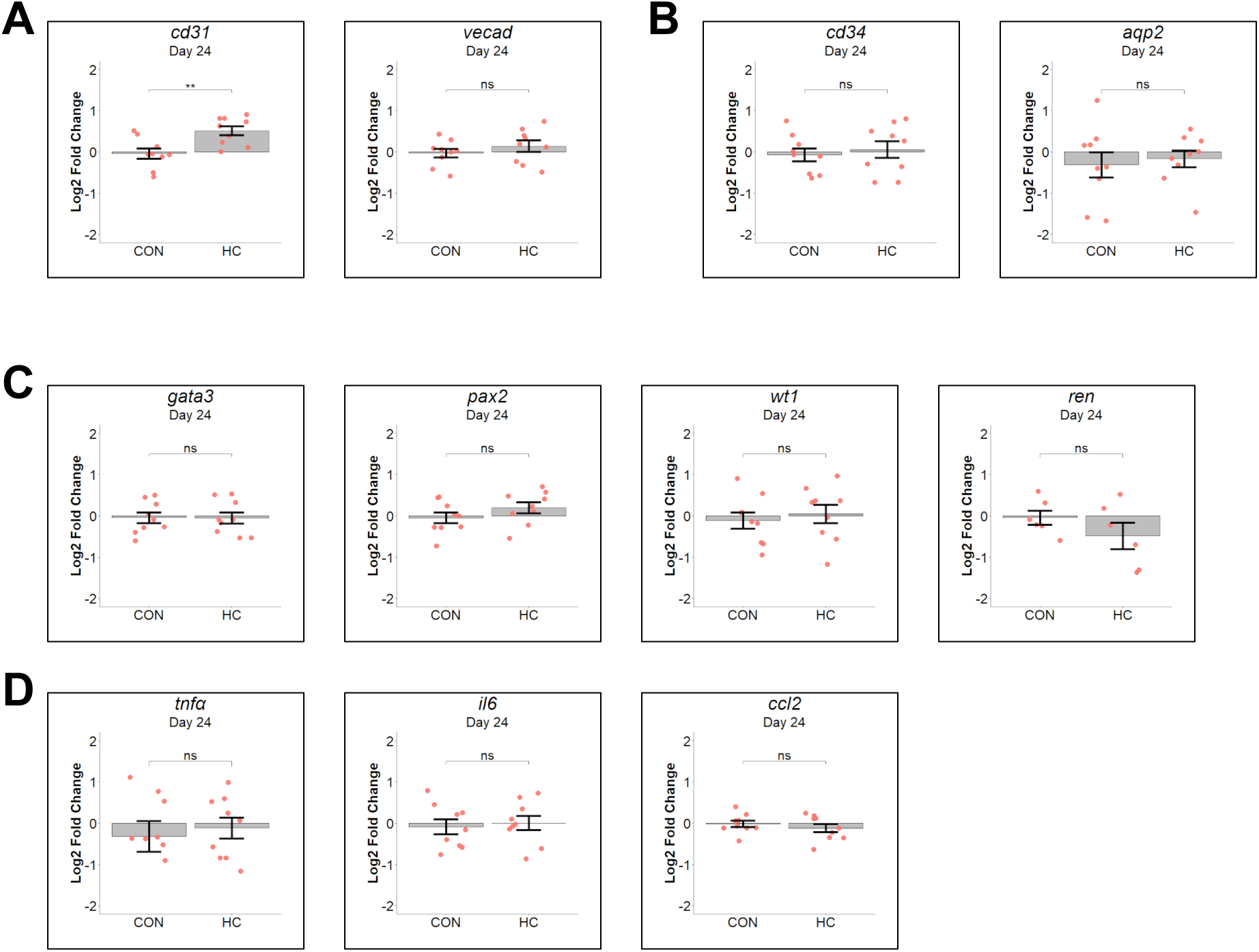
Characterization of canonical EC and kidney organoid gene expression on day 24 of differentiation. (A) Log2 fold change (HC-treated vs DMSO) in mRNA expression for EC junction genes *cd31* (left panel) and *vecad* (right panel). (B) Log2 fold change (HC-treated vs DMSO) in mRNA expression for hematopoietic progenitors (*cd34*; left panel) and collecting duct (*aqp2*; right panel). (C) Log2 fold change (HC-treated vs DMSO) in mRNA expression for ureteric bud (*gata3*; left panel), nephron progenitors (*pax2*; left-middle panel), total nephron (*wt1*; right-middle panel), and juxtaglomerular cells (*ren*; right panel). (D) Log2 fold change (HC-treated vs DMSO) in mRNA expression for pro-inflammatory cytokines *tnfa* (left panel), *il6* (middle panel), and *ccl2* (right panel). *n=9* biological replicates combined across 3 experiments per treatment group. Data are represented as mean ± SEM. * p< 0.05, ** p < 0.01, *** p < 0.001, and **** p < 0.0001.

**Supplemental Figure 2:**
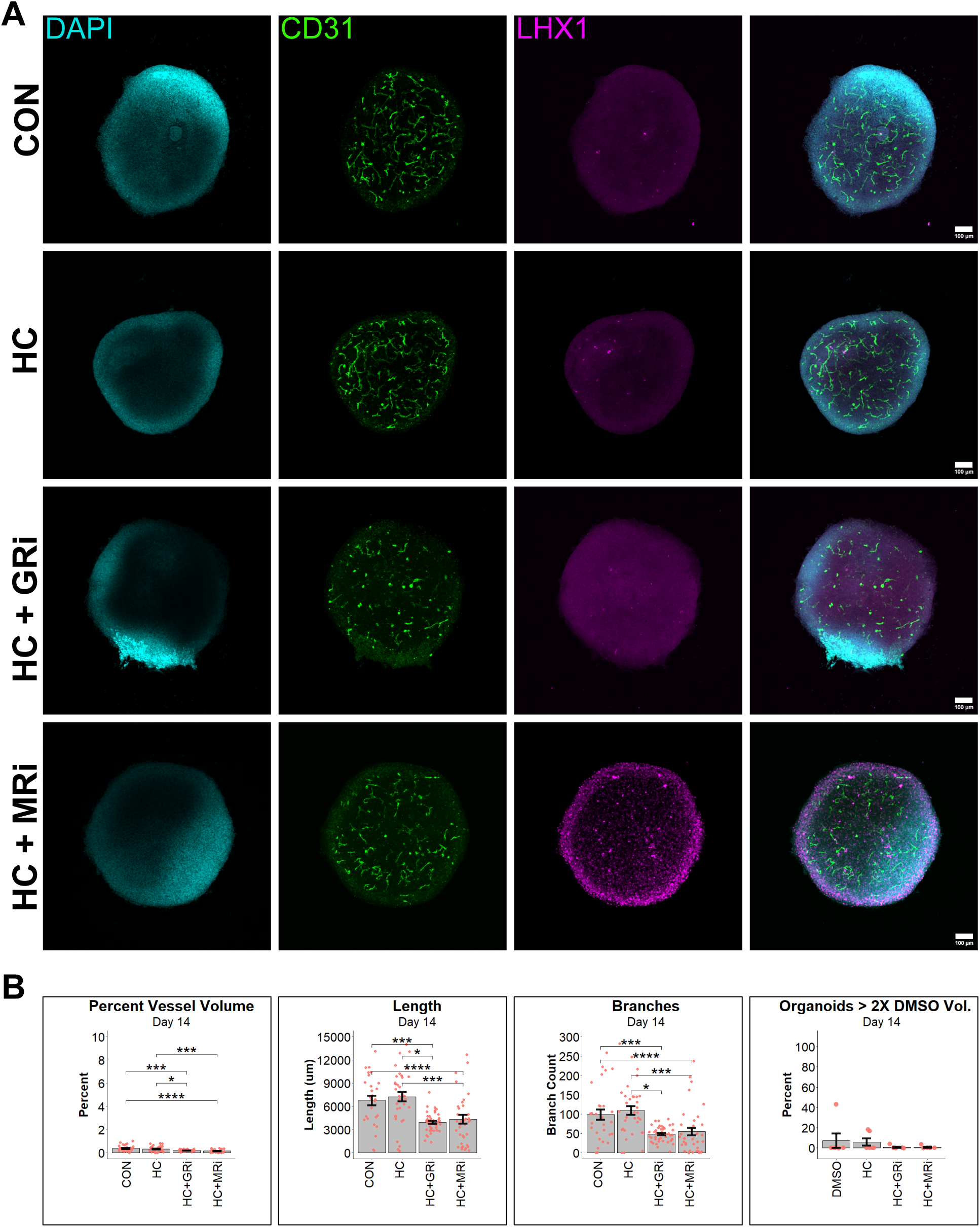
3D vessel analysis of GR and MR contribution to early vessel formation. (A) Representative images on day 14 of DAPI (cyan), CD31 (green), and LHX1 (magenta) in CON (1^st^ row), HC-(2^nd^ row), HC+GRi-(3^rd^ row), and HC+MRi-(4^th^ row) treated organoids. (B) Quantification of vessel volume as a percentage of the total organoid volume, per organoid (left panel), total vessel length per organoid (left-middle panel), total number of vessel branches per organoid (right-middle panel), and percentage of organoids with at least twice the mean percent vessel volume found in CON (right panel). Day 14 sample size (B): CON *n=28* organoids combined across 3 biological replicates, HC *n=34* organoids combined across 3 biological replicates, HC+GRi *n=43* organoids combined across 3 biological replicates, and HC+MRi *n=39* organoids combined across 3 biological replicates. Data are represented as mean ± SEM * p< 0.05, ** p < 0.01, *** p < 0.001, and **** p < 0.0001.

**Supplemental Figure 3:**
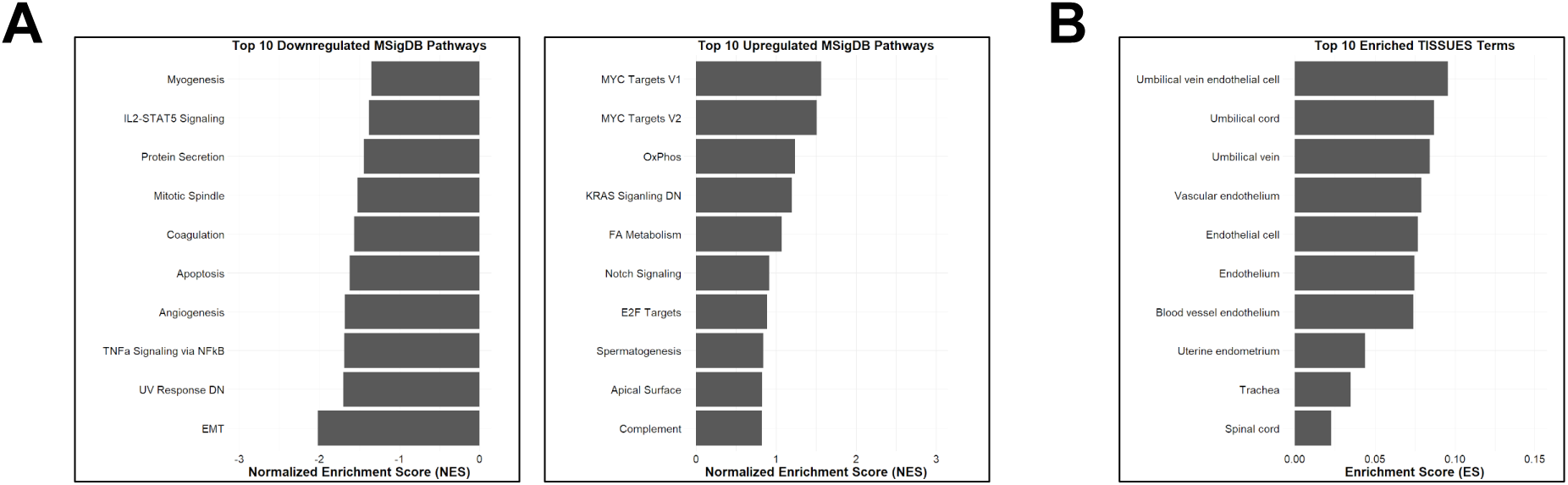
MSigDB Pathways and TISSUES terms analysis of day 9 organoids by bulk RNA-seq. (A) Top 10 downregulated MSigDB gene sets by normalized enrichment score (NES) and top 10 upregulated MSigDB genesets by NES. (B) Top 10 downregulated TISSUES terms.

**Supplemental Figure 4:**
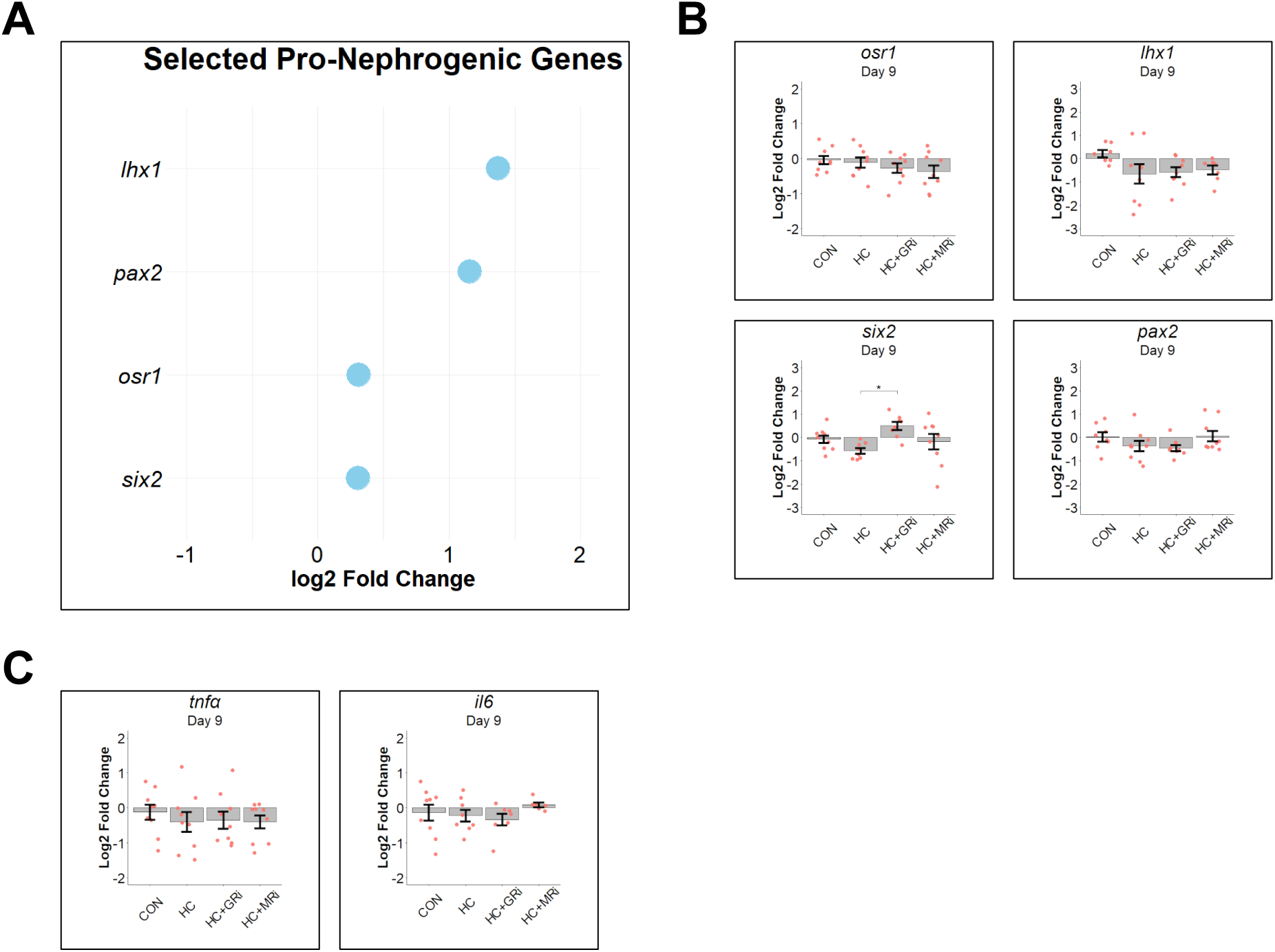
Analysis of selected pro-nephron and pro-inflammatory genes on day 9 of differentiation by bulk RNA-seq and RT-qPCR. (A) Dotplot of the fold change (HC-treated vs CON) in mRNA expression of selected canonical pro-nephrogenic genes identified in bulk RNA-seq. (B) Log2 fold change (HC-treated vs CON) in mRNA expression for mesoderm progenitors (*osr1*; top-left panel), nephron progenitors (*lhx1*; top-right panel, and *six2*; bottom-left panel), and tubular epithelial cells (*pax2*; bottom-right panel). (C) Log2 fold change (HC-treated vs CON) in mRNA expression for pro-inflammatory cytokines *tnfα* (left panel) and *il6* (right panel). *n=9* biological replicates combined across 3 experiments per treatment group. Data are represented as mean ± SEM. * p< 0.05, ** p < 0.01, *** p < 0.001, and **** p < 0.0001.

## References

Abdollahzadeh, F., Khoshdel-Rad, N., Bahrehbar, K., Erfanian, S., Ezzatizadeh, V., Totonchi, M., and Moghadasali, R. (2024). Enhancing maturity in 3D kidney micro-tissues through clonogenic cell combinations and endothelial integration. J Cell Mol Med 28, e18453. 10.1111/jcmm.18453.

Aceves, J.O., Heja, S., Kobayashi, K., Robinson, S.S., Miyoshi, T., Matsumoto, T., Schäffers, O.J.M., Morizane, R., and Lewis, J.A. (2022). 3D proximal tubule-on-chip model derived from kidney organoids with improved drug uptake. Sci Rep 12, 14997. 10.1038/s41598-022-19293-3.

Barrera-Chimal, J., and Jaisser, F. (2021). MR (Mineralocorticoid Receptor) in Endothelial Cells: A Major Contributor in Pulmonary Arterial Hypertension Remodeling. Hypertension 78, 466–468. 10.1161/hypertensionaha.121.17505.

Bridges, J.P., Sudha, P., Lipps, D., Wagner, A., Guo, M., Du, Y., Brown, K., Filuta, A., Kitzmiller, J., Stockman, C., et al. (2020). Glucocorticoid regulates mesenchymal cell differentiation required for perinatal lung morphogenesis and function. Am J Physiol Lung Cell Mol Physiol 319, L239–l255. 10.1152/ajplung.00459.2019.

Camarda, N.D., Lu, Q., Tesfu, A.F., Liu, R.R., Ibarrola, J., and Jaffe, I.Z. (2025). Mineralocorticoid Receptor in Endothelial Cells Contributes to Vascular Endothelial Growth Factor Receptor Inhibitor-Induced Vascular and Kidney Damage. Am J Hypertens 38, 104–110. 10.1093/ajh/hpae140.

Castanza, A.S., Recla, J.M., Eby, D., Thorvaldsdóttir, H., Bult, C.J., and Mesirov, J.P. (2023). Extending support for mouse data in the Molecular Signatures Database (MSigDB). Nature Methods 20, 1619–1620. 10.1038/s41592-023-02014-7.

Clerkin, S., Singh, K., Davis, J.L., Treacy, N.J., Krupa, I., Reynaud, E.G., Lees, R.M., Needham, S.R., MacWhite-Begg, D., Wychowaniec, J.K., et al. (2025). Tuneable gelatin methacryloyl (GelMA) hydrogels for the directed specification of renal cell types for hiPSC-derived kidney organoid maturation. Biomaterials 322, 123349. 10.1016/j.biomaterials.2025.123349.

Crossin, K.L., Tai, M.H., Krushel, L.A., Mauro, V.P., and Edelman, G.M. (1997). Glucocorticoid receptor pathways are involved in the inhibition of astrocyte proliferation. Proc Natl Acad Sci U S A 94, 2687–2692. 10.1073/pnas.94.6.2687.

Durant, S., Duval, D., and Homo-Delarche, F. (1986). Factors involved in the control of fibroblast proliferation by glucocorticoids: a review. Endocr Rev 7, 254–269. 10.1210/edrv-7-3-254.

Dyer, S.C., Austine-Orimoloye, O., Azov, A.G., Barba, M., Barnes, I., Barrera-Enriquez, V.P., Becker, A., Bennett, R., Beracochea, M., Berry, A., et al. (2024). Ensembl 2025. Nucleic Acids Research 53, D948–D957. 10.1093/nar/gkae1071.

Garreta, E., Moya-Rull, D., Marco, A., Amato, G., Ullate-Agote, A., Tarantino, C., Gallo, M., Esporrín-Ubieto, D., Centeno, A., Vilas-Zornoza, A., et al. (2024). Natural Hydrogels Support Kidney Organoid Generation and Promote In Vitro Angiogenesis. Adv Mater 36, e2400306. 10.1002/adma.202400306.

Gorini, S., Kim, S.K., Infante, M., Mammi, C., La Vignera, S., Fabbri, A., Jaffe, I.Z., and Caprio, M. (2019). Role of Aldosterone and Mineralocorticoid Receptor in Cardiovascular Aging. Front Endocrinol (Lausanne) 10, 584. 10.3389/fendo.2019.00584.

Grisé, K.N., Bautista, N.X., Jacques, K., Coles, B.L.K., and van der Kooy, D. (2021). Glucocorticoid agonists enhance retinal stem cell self-renewal and proliferation. Stem Cell Res Ther 12, 83. 10.1186/s13287-021-02136-9.

Huang, B., Zeng, Z., Kim, S., Fausto, C.C., Koppitch, K., Li, H., Li, Z., Chen, X., Guo, J., Zhang, C.C., et al. (2024). Long-term expandable mouse and human-induced nephron progenitor cells enable kidney organoid maturation and modeling of plasticity and disease. Cell Stem Cell 31, 921–939.e917. 10.1016/j.stem.2024.04.002.

Ibarrola, J., and Jaffe, I.Z. (2024). The Mineralocorticoid Receptor in the Vasculature: Friend or Foe? Annu Rev Physiol 86, 49–70. 10.1146/annurev-physiol-042022-015223.

Johnson, T.A., Fettweis, G., Wagh, K., Ceacero-Heras, D., Krishnamurthy, M., Sánchez de Medina, F., Martínez-Augustin, O., Upadhyaya, A., Hager, G.L., and Alvarez de la Rosa, D. (2024). The glucocorticoid receptor potentiates aldosterone-induced transcription by the mineralocorticoid receptor. Proceedings of the National Academy of Sciences 121, e2413737121. doi:10.1073/pnas.2413737121.

Kearney, H., Rak-Raszewska, A., Seijas-Gamardo, A., Escarda-Castro, E., Wieringa, P., Moroni, L., and Mota, C. (2025). Dimethyl sulfoxide primes induced pluripotent stem cells for more efficient nephron progenitor and kidney organoid differentiation. bioRxiv, 2025.2002.2007.637033. 10.1101/2025.02.07.637033.

Kim, J.W., Nam, S.A., Yi, J., Kim, J.Y., Lee, J.Y., Park, S.Y., Sen, T., Choi, Y.M., Lee, J.Y., Kim, H.L., et al. (2022). Kidney Decellularized Extracellular Matrix Enhanced the Vascularization and Maturation of Human Kidney Organoids. Adv Sci (Weinh) 9, e2103526. 10.1002/advs.202103526.

Kolde, R. (2019). Pheatmap: pretty heatmaps. R package version 1, 726.

Kroll, K.T., Homan, K.A., Uzel, S.G.M., Mata, M.M., Wolf, K.J., Rubins, J.E., and Lewis, J.A. (2024). A perfusable, vascularized kidney organoid-on-chip model. Biofabrication 16. 10.1088/1758-5090/ad5ac0.

Krupa, I., Treacy, N.J., Clerkin, S., Davis, J.L., Miller, A.F., Saiani, A., Wychowaniec, J.K., Reynaud, E.G., Brougham, D.F., and Crean, J. (2024). Protocol for the Growth and Maturation of hiPSC-Derived Kidney Organoids using Mechanically Defined Hydrogels. Curr Protoc 4, e1096. 10.1002/cpz1.1096.

Kuang, Z., Pang, C., Wang, H., Wei, X., Ye, X., Gao, X., and Sun, L. (2025). Generation of kidney organoids derived from human expanded potential stem cells. Cells Dev, 204025. 10.1016/j.cdev.2025.204025.

Kumar, S.V., Er, P.X., Lawlor, K.T., Motazedian, A., Scurr, M., Ghobrial, I., Combes, A.N., Zappia, L., Oshlack, A., Stanley, E.G., and Little, M.H. (2019). Kidney micro-organoids in suspension culture as a scalable source of human pluripotent stem cell-derived kidney cells. Development 146. 10.1242/dev.172361.

Liberzon, A., Birger, C., Thorvaldsdóttir, H., Ghandi, M., Mesirov, Jill P., and Tamayo, P. (2015). The Molecular Signatures Database Hallmark Gene Set Collection. Cell Systems 1, 417–425. 10.1016/j.cels.2015.12.004.

Lother, A., Deng, L., Huck, M., Fürst, D., Kowalski, J., Esser, J.S., Moser, M., Bode, C., and Hein, L. (2019). Endothelial cell mineralocorticoid receptors oppose VEGF-induced gene expression and angiogenesis. J Endocrinol 240, 15–26. 10.1530/joe-18-0494.

Love, M.I., Huber, W., and Anders, S. (2014). Moderated estimation of fold change and dispersion for RNA-seq data with DESeq2. Genome Biology 15, 550. 10.1186/s13059-014-0550-8.

Maggiore, J.C., LeGraw, R., Przepiorski, A., Velazquez, J., Chaney, C., Vanichapol, T., Streeter, E., Almuallim, Z., Oda, A., Chiba, T., et al. (2024). A genetically inducible endothelial niche enables vascularization of human kidney organoids with multilineage maturation and emergence of renin expressing cells. Kidney Int 106, 1086–1100. 10.1016/j.kint.2024.05.026.

Martinerie, L., Munier, M., Le Menuet, D., Meduri, G., Viengchareun, S., and Lombès, M. (2013). The mineralocorticoid signaling pathway throughout development: expression, regulation and pathophysiological implications. Biochimie 95, 148–157. 10.1016/j.biochi.2012.09.030.

McCann, K.E., Lustberg, D.J., Shaughnessy, E.K., Carstens, K.E., Farris, S., Alexander, G.M., Radzicki, D., Zhao, M., and Dudek, S.M. (2021). Novel role for mineralocorticoid receptors in control of a neuronal phenotype. Molecular Psychiatry 26, 350–364. 10.1038/s41380-019-0598-7.

Morizane, R., Lam, A.Q., Freedman, B.S., Kishi, S., Valerius, M.T., and Bonventre, J.V. (2015). Nephron organoids derived from human pluripotent stem cells model kidney development and injury. Nat Biotechnol 33, 1193–1200. 10.1038/nbt.3392.

Nerger, B.A., Sinha, S., Lee, N.N., Cheriyan, M., Bertsch, P., Johnson, C.P., Mahadevan, L., Bonventre, J.V., and Mooney, D.J. (2024). 3D Hydrogel Encapsulation Regulates Nephrogenesis in Kidney Organoids. Adv Mater 36, e2308325. 10.1002/adma.202308325.

Nesan, D., Kamkar, M., Burrows, J., Scott, I.C., Marsden, M., and Vijayan, M.M. (2012). Glucocorticoid receptor signaling is essential for mesoderm formation and muscle development in zebrafish. Endocrinology 153, 1288–1300. 10.1210/en.2011-1559.

Palasca, O., Santos, A., Stolte, C., Gorodkin, J., and Jensen, L.J. (2018). TISSUES 2.0: an integrative web resource on mammalian tissue expression. Database 2018. 10.1093/database/bay003.

Perens, E.A., Garavito-Aguilar, Z.V., Guio-Vega, G.P., Peña, K.T., Schindler, Y.L., and Yelon, D. (2016). Hand2 inhibits kidney specification while promoting vein formation within the posterior mesoderm. eLife 5, e19941. 10.7554/eLife.19941.

Przybyciński, J., Drożdżal, S., Domański, L., Dziedziejko, V., and Pawlik, A. (2021). Role of Endothelial Glucocorticoid Receptor in the Pathogenesis of Kidney Diseases. Int J Mol Sci 22. 10.3390/ijms222413295.

Ribatti, D., Ligresti, G., and Nicosia, R.F. (2023). Kidney endothelial cell heterogeneity, angiocrine activity and paracrine regulatory mechanisms. Vascul Pharmacol 148, 107139. 10.1016/j.vph.2022.107139.

Roberts, E.W., Denton, A.E., and Fearon, D.T. (2016). Roles of Stromal Cells in the Immune System. In Encyclopedia of Cell Biology (Second Edition), R.A. Bradshaw, G.W. Hart, and P.D. Stahl, eds. (Academic Press), pp. 484–492. 10.1016/B978-0-12-821618-7.30079-7.

Ruiter, F.A.A., Morgan, F.L.C., Roumans, N., Schumacher, A., Slaats, G.G., Moroni, L., LaPointe, V.L.S., and Baker, M.B. (2022). Soft, Dynamic Hydrogel Confinement Improves Kidney Organoid Lumen Morphology and Reduces Epithelial-Mesenchymal Transition in Culture. Adv Sci (Weinh) 9, e2200543. 10.1002/advs.202200543.

Salvador, E., Shityakov, S., and Förster, C. (2014). Glucocorticoids and endothelial cell barrier function. Cell Tissue Res 355, 597–605. 10.1007/s00441-013-1762-z.

Sarami, I., Hekman, K.E., Gupta, A.K., Snider, J.M., Ivancic, D., Zec, M., Kandpal, M., Ben-Sahra, I., Menon, R., Otto, E.A., et al. (2025). Parallel multiOMIC analysis reveals glutamine deprivation enhances directed differentiation of renal organoids. bioRxiv. 10.1101/2025.02.27.640060.

Srivastava, S., Nataraj, N.B., Sekar, A., Ghosh, S., Bornstein, C., Drago-Garcia, D., Roth, L., Romaniello, D., Marrocco, I., David, E., et al. (2019). ETS Proteins Bind with Glucocorticoid Receptors: Relevance for Treatment of Ewing Sarcoma. Cell Rep 29, 104–117.e104. 10.1016/j.celrep.2019.08.088.

Srivastava, S.P., Zhou, H., Setia, O., Liu, B., Kanasaki, K., Koya, D., Dardik, A., Fernandez-Hernando, C., and Goodwin, J. (2021). Loss of endothelial glucocorticoid receptor accelerates diabetic nephropathy. Nat Commun 12, 2368. 10.1038/s41467-021-22617-y.

Subramanian, A., Tamayo, P., Mootha, V.K., Mukherjee, S., Ebert, B.L., Gillette, M.A., Paulovich, A., Pomeroy, S.L., Golub, T.R., Lander, E.S., and Mesirov, J.P. (2005). Gene set enrichment analysis: A knowledge-based approach for interpreting genome-wide expression profiles. Proceedings of the National Academy of Sciences 102, 15545–15550. doi:10.1073/pnas.0506580102.

Szklarczyk, D., Kirsch, R., Koutrouli, M., Nastou, K., Mehryary, F., Hachilif, R., Gable, A.L., Fang, T., Doncheva, N.T., Pyysalo, S., et al. (2023). The STRING database in 2023: protein-protein association networks and functional enrichment analyses for any sequenced genome of interest. Nucleic Acids Res 51, D638–d646. 10.1093/nar/gkac1000.

Vanslambrouck, J.M., Tan, K.S., Mah, S., and Little, M.H. (2023). Generation of proximal tubule-enhanced kidney organoids from human pluripotent stem cells. Nat Protoc 18, 3229–3252. 10.1038/s41596-023-00880-1.

Vegiopoulos, A., and Herzig, S. (2007). Glucocorticoids, metabolism and metabolic diseases. Mol Cell Endocrinol 275, 43–61. 10.1016/j.mce.2007.05.015.

Wang, J., Chen, F., Zhu, S., Li, X., Shi, W., Dai, Z., Hao, L., and Wang, X. (2022). Adverse effects of prenatal dexamethasone exposure on fetal development. Journal of Reproductive Immunology 151, 103619. 10.1016/j.jri.2022.103619.

Wickham, H., Chang, W., and Wickham, M.H. (2016). Package ‘ggplot2’. Create elegant data visualisations using the grammar of graphics. Version 2, 1–189.

Wilson, S.B., Santos, I.P., Wildfang, L., Imsa, K., and Little, M.H. (2025). Generation of multi-lineage kidney assembloids with integration between nephrons and a single exiting collecting duct. bioRxiv, 2025.2002.2027.640561. 10.1101/2025.02.27.640561.

Young, M.J., and Clyne, C.D. (2021). Mineralocorticoid receptor actions in cardiovascular development and disease. Essays Biochem 65, 901–911. 10.1042/ebc20210006.

Zielińska, K.A., Van Moortel, L., Opdenakker, G., De Bosscher, K., and Van den Steen, P.E. (2016). Endothelial Response to Glucocorticoids in Inflammatory Diseases. Front Immunol 7, 592. 10.3389/fimmu.2016.00592.

