## Supplemental File 1 for "Glucocorticoid and Mineralocorticoid Receptors Jointly Promote Vascular Development in Kidney Organoids": nf_core_rnaseq_software_mqc_versions.yml

```
BEDTOOLS_GENOMECOV_FW:
  bedtools: 2.31.1
CUSTOM_GETCHROMSIZES:
  getchromsizes: 1.21
CUSTOM_TX2GENE:
  python: 3.10.4
DESEQ2_QC_STAR_SALMON:
  r-base: 4.0.3
  bioconductor-deseq2: 1.28.0
DUPRADAR:
  bioconductor-dupradar: 1.32.0
FASTQC:
  fastqc: 0.12.1
FQ_LINT:
  fq: 0.12.0 (2024-07-08)
FQ_SUBSAMPLE:
  fq: 0.12.0 (2024-07-08)
GTF2BED:
  perl: 5.26.2
GTF_FILTER:
  python: 3.9.5
GUNZIP_FASTA:
  gunzip: 1.1
GUNZIP_GTF:
  gunzip: 1.1
MAKE_TRANSCRIPTS_FASTA:
  rsem: 1.3.1
  star: 2.7.10a
MULTIQC_CUSTOM_BIOTYPE:
  python: 3.9.5
PICARD_MARKDUPLICATES:
  picard: 3.1.1
RSEQC_BAMSTAT:
  rseqc: 5.0.2
RSEQC_INFEREXPERIMENT:
  rseqc: 5.0.2
RSEQC_INNERDISTANCE:
  rseqc: 5.0.2
RSEQC_JUNCTIONANNOTATION:
  rseqc: 5.0.2
RSEQC_JUNCTIONSATURATION:
  rseqc: 5.0.2
RSEQC_READDISTRIBUTION:
  rseqc: 5.0.2
RSEQC_READDUPLICATION:
  rseqc: 5.0.2
SALMON_INDEX:
  salmon: 1.10.3
SALMON_QUANT:
  salmon: 1.10.3
SAMTOOLS_FLAGSTAT:
  samtools: 1.21
SAMTOOLS_IDXSTATS:
  samtools: 1.21
SAMTOOLS_INDEX:
  samtools: 1.21
SAMTOOLS_SORT:
  samtools: 1.21
SAMTOOLS_STATS:
  samtools: 1.21
SE_GENE:
  bioconductor-summarizedexperiment: 1.32.0
STAR_ALIGN:
  star: 2.7.11b
  samtools: 1.21
  gawk: 5.1.0
STAR_GENOMEGENERATE:
  star: 2.7.11b
  samtools: 1.21
  gawk: 5.1.0
STRINGTIE_STRINGTIE:
  stringtie: 2.2.3
SUBREAD_FEATURECOUNTS:
  subread: 2.0.6
TRIMGALORE:
  trimgalore: 0.6.10
  cutadapt: 4.9
TXIMETA_TXIMPORT:
  bioconductor-tximeta: 1.20.1
UCSC_BEDCLIP:
  ucsc: 377
UCSC_BEDGRAPHTOBIGWIG:
  ucsc: 469
Workflow:
    nf-core/rnaseq: v3.18.0-gb96a753
    Nextflow: 24.04.4
```
